# Outcomes of Progranulin Gene Therapy in the Retina are Dependent on Time of Delivery

**DOI:** 10.1101/2021.02.24.432570

**Authors:** Emilia A. Zin, Daisy Han, Jennifer Tran, Nikolas Morisson-Welch, Meike Visel, Mervi Kuronen, John G. Flannery

## Abstract

Neuronal ceroid lipofuscinosis (NCL) is a family of neurodegenerative diseases caused by mutations to genes related to lysosomal function. One variant, CNL11, is caused by mutations to the gene encoding the protein progranulin. Primarily secreted by microglia, progranulin regulates neuronal lysosomal function once endocytosed. Absence of progranulin causes cerebellar atrophy, seizures, ataxia, dementia and vision loss. As progranulin gene therapies targeting the brain are developed, it is also advantageous to focus on the retina, as its characteristics are beneficial for gene therapy development: the retina is easily visible through direct imaging, can be assessed through quantitative methods *in vivo*, requires smaller amounts of AAV and AAV can be administered via a less invasive surgery. In this study we characterize the retinal degeneration in a progranulin knockout mouse model of CLN11 and study the effects of gene replacement at different time points. All mice heterologously expressing progranulin showed reduction in lipofuscin deposits and microglia infiltration. While mice that receive systemic AAV9.2YF-scCAG-PGRN at post-natal day 3 or 4 show a reduction in retina thinning, mice injected intravitreally at months 1 and 6 with 7m8-scCAG-PGRN show no improvement, and mice injected at 12 months of age show increased retinal thinning in comparison to their controls. Thus, delivery of progranulin proves to be time-sensitive, requiring early administration for optimal therapeutic benefit.

## Introduction

Neuronal ceroid lipofuscinosis (NCL), or Batten disease, is a family of disorders characterized by the accumulation of autofluorescent lipofuscin in neurons and clinically, with seizures, progressive vision loss, and motor and cognitive decline(1). There are at least 14 different variants of NCL, each caused by mutations to different genes. These variants are labeled sequentially: CLN1, CLN2, CLN3, and so forth(2, 3). CLN11, identified in 2012 in a pair of Italian siblings, is caused by biallelic mutations in the progranulin *GRN* (or *PGRN*) gene(4). These patients present vision loss, seizures, ataxia and cognitive decline(5). Interestingly, single allele mutations in *GRN* are also one of the causes of frontotemporal dementia (FTD)(6, 7).

Progranulin is an 88 kDa glycoprotein expressed in multiple cell types(8). The full protein is composed of seven and a half tandem repeats of cysteine-rich sequences. Progranulin is mainly secreted by microglia and endocytosed by neurons through sortilin receptors, then shuttled to the endolysosome and early lysosome, where it is cleaved into the seven separate granulins by three different lysosomal proteases: cathepsins B, D and L(9–11).

Progranulin is known to directly or indirectly affect lysosomal biogenesis and autophagy through transcriptional regulation. Transcriptional activation of lysosomal genes through the CLEAR sequence, and consequential lysosomal upregulation, seem to be related to protein aggregation and mislocalization(12). This leads to one of the hallmarks of NCL: accumulation of lipofuscin, an autofluorescent granule deposit composed of proteins, lipids, sugars and metals(13). Lipofuscin naturally accumulates in several tissues with age, but mitochondria, membrane or lysosomal damage can increase lipofuscin deposits. Since all NCLs present with some form of lysosomal damage, accumulation of lipofuscin represents a hallmark of the disease, which also applies to CLN11(14). As such, lipofuscin accumulation can be used as a biomarker for NCL therapies(15).

Most types of NCL are exhibited by animal models that replicate disease progression. For CLN11 there are several mouse models in which *GRN* was knocked out by different methods(16, 17). The homozygous PGRN^-/-^ mouse line used in this study shows slow neurodegeneration, with loss of neurons by 12 months of age, increase in microglial activation, mild behavioral alterations, but no seizures(17, 18). Several studies have analyzed neuronal degeneration in the mouse brain, but very few have reported on retinal degeneration progression, which is addressed in this study (19–21).

Gene replacement represents an excellent strategy for NCL therapy; almost all NCLs are autosomal recessive, so the delivery of a functional copy of the mutated gene should theoretically restore cellular function. There are several promising NCL gene therapies currently under preclinical and clinical development(22, 23). There are numerous preclinical mouse studies, including a gene therapy for CLN8, where expression of the *CLN8* gene, delivered via an AAV9 capsid by an intracerebroventricular injection, prolongs the mutant’s lifespan and restores normal behavior(24). Similarly, CLN6^nclf^ mice, which show photoreceptor degeneration, had reduced cell death after the intravitreal delivery of *CLN6* to bipolar cells through the 7m8 capsid(25, 26). Two studies have tackled gene therapy in PGRN^-/-^ mice: one reports improvement in lysosomal function after bilateral stereotaxic injections of rAAV2-CBA-PGRN in the medial prefrontal cortex, and the other describes hippocampal atrophy due to T-cell toxicity after unilateral stereotaxic posterior right lateral ventricle injections of rAAV4-CAG-PGRN or rAAV9-CMV-PGRN(27, 28). However, none looked at effects of PGRN gene therapy in the retina.

A retinal focused study could provide critical insights into the effects of a progranulin gene therapy. Not only is the retina a window to the brain, it is also easily monitored over time through electrophysiology and direct imaging. Furthermore, most patients with biallelic *GRN* mutations show vision loss, and would likely require viral delivery targeted to the retina itself. Given the already published conflicting results and a GRN-FTD clinical trial underway, a gene therapy study targeting progranulin to the retina would provide further data on treatment efficacy over time(29).

We hypothesized that by delivering PGRN to all cell layers of the retina via AAV vectors, enough progranulin protein would be produced intracellularly and secreted to the extracellular space to prevent neuronal death, with the assumption that early delivery might completely stall retinal degeneration. For such purpose, the AAV2.7m8 and AAV9.2YF capsids were chosen. 7m8 was selected as an engineered capsid capable of reaching all cell layers in the retina when injected from the vitreous in adult mice(30). As AAV9 can cross the blood-retina and blood-brain barriers in mice until 7 days old, it seemed an appropriate choice for intravenous deliveries in mice pups between 3 and 4 days of age, where direct retinal delivery is challenging due to eye size and closed eyelids(31). The included mutations on the AAV9 capsid, i.e., Y446F and Y731F (AAV9.2YF) improve gene transfer by inhibiting the proteasomal degradation of the AAV capsid. In both constructs, the ubiquitous and synthetic promoter CAG was used in a self-complementary (sc) plasmid. Animals received intravitreal injections with 7m8-scCAG-mPGRN at 1 month, 6 months and 12 months of age. Mice that received a tail vein injection at post-natal day 3 or 4 had AAV92YF-scCAG-mPGRN delivered.

## Methods

### Production of Viral Vectors

rAAV was packaged in HEK293T cells by transient plasmid co-transfection as previously described(32). The capsid serotype plasmids were AAV2.7m8 (7m8) or AAV9.2YF Rep/Cap, scCAG-mPGRN or scCAG-hPGRN were used as the transgene, and an adenovirus (Ad) Helper plasmid(32) was used in all preps. The virus was purified with iodixanol ultracentrifugation gradient, and buffers exchanged with Amicon Ultra-15 Centrifugal Filter Units, so the final viral stock was in sterile PBS. Viral titers were established via qPCR, with primers for ITRs.

### Animals and Injections

The experiments were authorized by the Animal Care and Use Committee, Office of Animal Care and Use at UC Berkeley (AUP 2014-09-6705). All experiments were conducted according to the ARVO Statement for the Use of Animals and the guidelines of the Office of Laboratory Animal Care at the University of California, Berkeley, CA. PGRN^+/-^ mice were obtained from the Gladstone Institute at UCSF, as originally described by Martens et al, 2012(18). Initially PGRN^+/-^ mice were backcrossed into C57BL/6J and then PGRN^-/-^ breeding pairs were set up. All mice were housed at a 12h light/dark cycle with *ad libitum* access to food and water. For intravitreal injections, mice were anesthetized with intraperitoneal injections of ketamine (72 mg/kg) and xylazine (64 mg/kg). To gain entry into the vitreous chamber, the sclera was punctured posterior to the limbus with a disposable 30 ½ gauge needle. A blunt needle attached to a glass Hamilton syringe was used to deliver 2 μl of AAV via the same path into the vitreous cavity, over the optic nerve head. Contralateral control eyes were injected with 2 μl of sterile PBS. Intravenous injections in pups were given via the tail vein using sterile BD Ultra Fine (31 gauge) Insulin syringes. Corresponding injected viral titers can be found in Table 1.

**Table 1.** AAV viral titers established through qPCR.

| **rAAV (100-150 μl)** | **Titer (vg/ml)** |
| --- | --- |
| 7m8-scCAG-mPGRN | 3.73E+13 |
| 7m8-scCAG-mPGRN | 3.17E+14 |
| 7m8-scCAG-hPGRN | 3.0E+12 |
| 7m8-scCAG-hPGRN | 1.84E+13 |
| AAV92YF-scCAG-mPGRN | 1.20E+14 |
| AAV92YF-scCAG-hPGRN | 1.10E+14 |

### Electroretinogram (ERG) Recording

Mice were dark adapted overnight in dark boxes, and ERGs performed between 8:30 am and 12:30 pm. Mice were anesthetized with intraperitoneal injections of ketamine and xylazine as aforementioned, and one drop of tropicamide (1%) and phenylephrine (2.5%) each applied topically for pupil dilation. Mice were then positioned on a heated platform and contact lens recording electrodes placed on corneas of both eyes, with a ground electrode inserted subcutaneously between both eyes, and a reference electrode introduced subcutaneously to the tail. Scotopic ERGs were firstly recorded as three flashes at 1 log cd x s/m^2^ on a dark background and averaged. Rod responses were then saturated by a bright background for 5 minutes. Secondly, photopic responses were recorded at 1.4 log cd x s/m^2^ and presented on a light background. ERGs were recorded with an Espion E2 system (Diagnosys, LLC, Lowell, MA) and data was analyzed with MatLab (MathWorks, Natick, MA) to extract A and B wave amplitudes from recordings.

### Optical Coherence Tomography (OCT)

Retinal images were captured *in vivo* using an 840 nm SDOIS OCT system (Bioptigen, Durham, NC) including an 840 nm SDOIS Engine with 93 nm bandwidth internal source providing a tissue resolution of less than 3.0 μm. Animals were anesthetized with intraperitoneal injections of ketamine and xylazine, and pupils dilated with tropicamide and phenylephrine, as previously described. Superior and inferior retinal hemispheres were imaged in 96 sections, which were averaged into 8 slices. The thickness of the total retina, from inner limiting membrane (ILM) to the retinal pigment epithelium (RPE), and photoreceptor layers, from outer nuclear layer (ONL) to RPE, were measured with InVivoVue software (Bioptigen, Durham, NC) and FIJI (Fiji is Just ImageJ, National Institute of Health, Bethesda, MD)(33) from the superior and inferior sections that showed the optic nerve head.

### Immunohistochemistry

Mice were euthanized with CO2 overdose and cervical dislocation. Eyes were enucleated and fixed in paraformaldehyde 4% overnight, then washed in PBS. Eyes were dissected, cornea, lens and vitreous removed, and the remaining eye cups incubated in 30% sucrose overnight. Eye cups were then imbedded in optimal cutting temperature compound (OCT), and sectioned with a Leica CM3050 cryostat (Leica Biosystems, Buffalo Grove, IL). Retinal sections were washed with PBS, incubated with Triton X-100 0.5% and BSA 1% for 15 minutes, then BSA 1% for 1 hour. Sections were incubated overnight with primary antibodies (Table 2) in BSA 1%. On the following day slides were washed three times with PBS, then incubated with secondary antibodies (Table 2) in BSA 1% for 1 hour at room temperature. After two PBS washes, sections were mounted with Vectashield Antifade Mounting Medium with DAPI. Slides were imaged with a laser scanning confocal microscope (LSM710, Carl Zeiss, Oberkochen, Germany).

**Table 2.** Antibodies and their dilutions.

| Antibody | Company | Dilution |
| --- | --- | --- |
| Sheep anti-mPGRN | R&D Systems (Minneapolis, MN), AF2557 | 1:500 |
| Goat anti-hPGRN | R&D Systems, AF2420 | 1:500 |
| Rabbit anti-IBA1 | Fujifilm Wako Diagnostics (Mountain View, CA), 019-19741 | 1:1000 |
| Rabbit anti-LAMP1 | Abcam (Cambridge, UK), ab24170 | 1:1000 |
| Alexa Fluor 488 anti-goat | ThermoFisher (Waltham, MA), A32814 | 1:2000 |
| Alexa Fluor 594 anti-sheep | ThermoFisher, A11016 | 1:2000 |
| Alexa Fluor 594 anti-rabbit | ThermoFisher, A21207 | 1:2000 |

### Lipofuscin Imaging and Quantification

After sectioning, retinas were incubated with Triton X-100 0.5% and BSA 1% for 15 minutes, washed with PBS and mounted with Vectashield Antifade Mounting Medium and DAPI. All sections used for lipofuscin quantification were centered on the optic nerve. Sections were imaged with either 488 nm-laser or 564 nm-laser with the same settings. Images were processed with FIJI, then transformed into binary images and particles were counted. Total area covered by autofluorescent particles was divided by the total area covered by DAPI, and multiplied by 100.

## Results

PGRN^-/-^ mice have been used in several studies to understand development of NCL and FTD, and although pathological degeneration in the brain has been studied extensively, the timeline of retinal development is not as well understood. Previous studies reported retinal thinning by 12 and 18 months of age but did not report on earlier timepoints(20). Therefore, retinas of PGRN^-/-^ and C57BL/6J mice were compared over time (Figure 1). To explore retinal thinning, overall retinal (inner-limiting membrane to retinal pigment epithelium) and photoreceptor layer (outer nuclear layer to retinal pigment epithelium) thickness were measured by optical coherence tomography (OCT) imaging at month 1, month 5, month 12 and month 18 of age (Figure 1A, B, C, D, respectively). PGRN^-/-^ photoreceptor layers were already significantly thinner than those of C57BL/6J retinas at 1 month of age (Figure 1A), and this trend continued at 5, 12 and 18 months of age (Figure 1B, C, D). However, statistically significant overall retinal thinning of PGRN^-/-^ retinas only occurred at month 12 and 18 (Figure 1C, D). Differences varied between 26.38 μm and 16.64 μm for total retinas at 12 months, and between 35.80 μm and 26.33 μm for total retinas at 18 months old.

**Figure 1.**
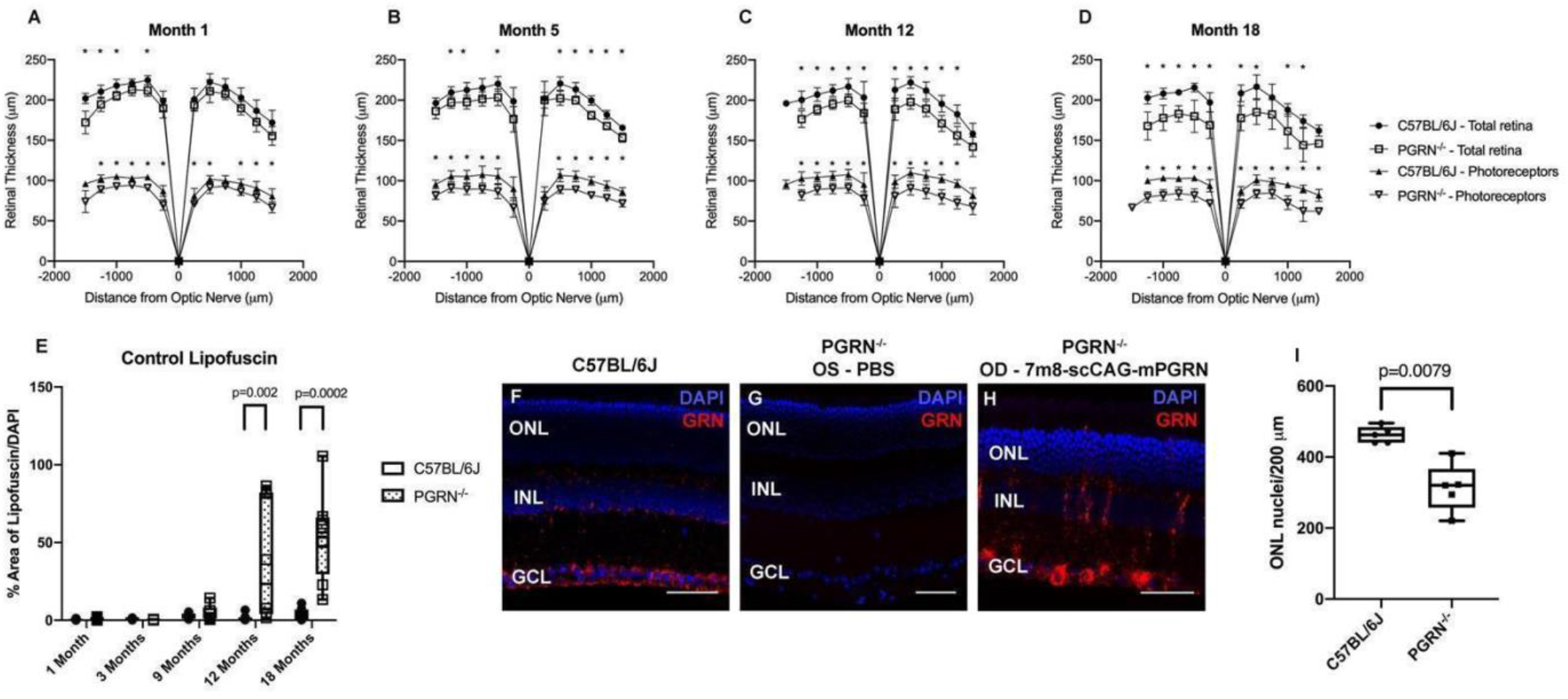
Retinal degeneration in PGRN^-/-^ mice over time. Optical coherence tomography (OCT) measurements in superior (−250 to −1500 μm from the optic nerve) and inferior (250 to 1500 μm from optic nerve) retinas. Total retina thickness was measured from the ILM to RPE (upper curves: C57BL/6J in closed circles and PGRN^-/-^ in open squares), and photoreceptor layer thickness measured from ONL to RPE (lower curves: C57BL/6J in closed triangles and PGRN^-/-^ in open triangles). Month 1 (A), month 5 (B), month 12 (C) and month 18 (D) were recorded. Error bars represent ± standard deviation, and closed circles or triangles stand for mean values for retinal thickness of N=7 animals per group for C57BL/6J at months 1 and 18, N=6 for month 5 and N=9 for month 12 animals per group. Open squares or triangles stand for mean values for retinal thickness for PGRN^-/-^ mice of N=5 animals per group at months 1, 5 and 18, and N=10 for month 12. Statistical tests were performed at each distance from optic nerve between mouse strains (t-tests), asterisks representing p < 0.05. Lipofuscin was mainly found in PGRN^-/-^ retinas and quantified by measuring total area covered by lipofuscin in flatmounts over total area covered by nuclei in DAPI at different time points (E). Box and whiskers plot, boxes show interquartile range (25% to 75%), whiskers represent minimum and maximum values, horizontal lines represent median values of 6 animals at month 3 and month 9, and 8 animals at month 1, 12 and 18. Mann-Whitney statistical test between mouse strains at different time points, p values as indicated, N=6 or N=8 per group. Immunohistochemistry for PGRN in either C57BL/6J retina (F), PGRN^-/-^ OS retina with PBS injection (G), or PGRN^-/-^ OD retina expressing mouse PGRN through 7m8-scCAG-mPGRN intravitreal injection (H). DAPI staining nuclei (blue) and PGRN (red). Scale bars: 50 μm. Quantification of photoreceptor nuclei in C57BL/6J and PGRN^-/-^ retinas, where nuclei in outer nuclear layer were quantified along 200 μm (I). Box and whiskers plot, boxes show interquartile range (25% to 75%), whiskers represent minimum and maximum values, horizontal lines represent median values of 5 animals/group. Inter-strain difference was statistically significant (Mann-Whitney test, p=0.0079). Scale bars = 50 μm

ERGs reflect the electrical activity of retinal neurons and glial cells generated in response to light. A-waves reflect photoreceptor activity and B waves, ON-bipolar and Müller glia activity. Cone and rod responses can be measured by photopic or scotopic ERGs, respectively. A- and B-wave amplitude data derived from scotopic and photopic ERGs from C57BL/6J and PGRN^-/-^ mice at 1, 3, 7, 9 and 12 months of age are shown in Supplemental Figure 1. Significant strain-dependent differences in scotopic A wave amplitudes were obtained at 7, 9 and 12 months of age (Sup Figure 1A).

A previous study by Ward and colleagues report increases in retinal lipofuscinosis at months 1.5, 3, 6, 12, and 18(34). To replicate their findings, retinal flatmounts were imaged, and lipofuscin was visible principally in PGRN^-/-^ retinas (Figure 1E). PGRN^-/-^ retinas showed a trend in lipofuscin increase when compared to C57BL/6J retinas, and the difference became statistically significant at months 12 and 18. Hence, while animal to animal variation must be taken into consideration for PGRN^-/-^ mice, these results can support the use of retinal lipofuscin content as a quantifiable biomarker for this retinal disease as well as treatment efficacy.

Endogenous expression of PGRN in C57BL/6J mice was compared to expression driven by 7m8-scCAG-mPGRN (Figure 1F, G, H). The AAV construct was intravitreally injected in the right eye (OD) of PGRN^-/-^ mice, and PBS in the left eye (OS). Endogenous PGRN expression was mainly seen in the ganglion cell layer (GCL) and inner nuclear layer (INL), with some expression in the ONL (Figure 1F). Meanwhile, as expected, PGRN^-/-^ eyes injected with PBS showed no PGRN expression (Figure 1G), while retinas injected with 7m8-scCAG-mPGRN transduced mouse PGRN in GCL, INL, Müller glia and occasional photoreceptors (Figure 1H), confirming that PGRN^-/-^ retinas could indeed widely express PGRN through a ubiquitous promoter 8 months post-injection.

Mice intravitreally injected at 1 and 6 months old with either 7m8-scCAG-mPGRN or 7m8-scCAG-hPGRN were euthanized at 12-months-old, enucleated and retinal tissue collected for processing. Immunohistochemistry for mouse PGRN (mPGRN) and human PGRN (hPGRN) showed an overall increase in PGRN expression in PGRN^-/-^ mice injected at 6 months of age (Figure 2). PGRN was expressed mainly in the GCL (Figure 2A, E, I), but can also be found in photoreceptors and Müller glia (Figure 2A), and bipolar cells (Figure 2I). Both animals F2 and F1 were of the same litter, injected with the same viral titer (3.97E+13 vg/ml) and volume (2 μl), but demonstrated how gene transduction can vary from animal to animal, with higher progranulin protein expression seen in Figure 2A. Interestingly, microglia infiltrated the INL in all panels (arrowheads, Figure 2D, G, H, K, L), but not in Figure 2C, showing infiltration in all contralateral control retinas, but not in one retina expressing PGRN (Figure 2C).

**Figure 2.**
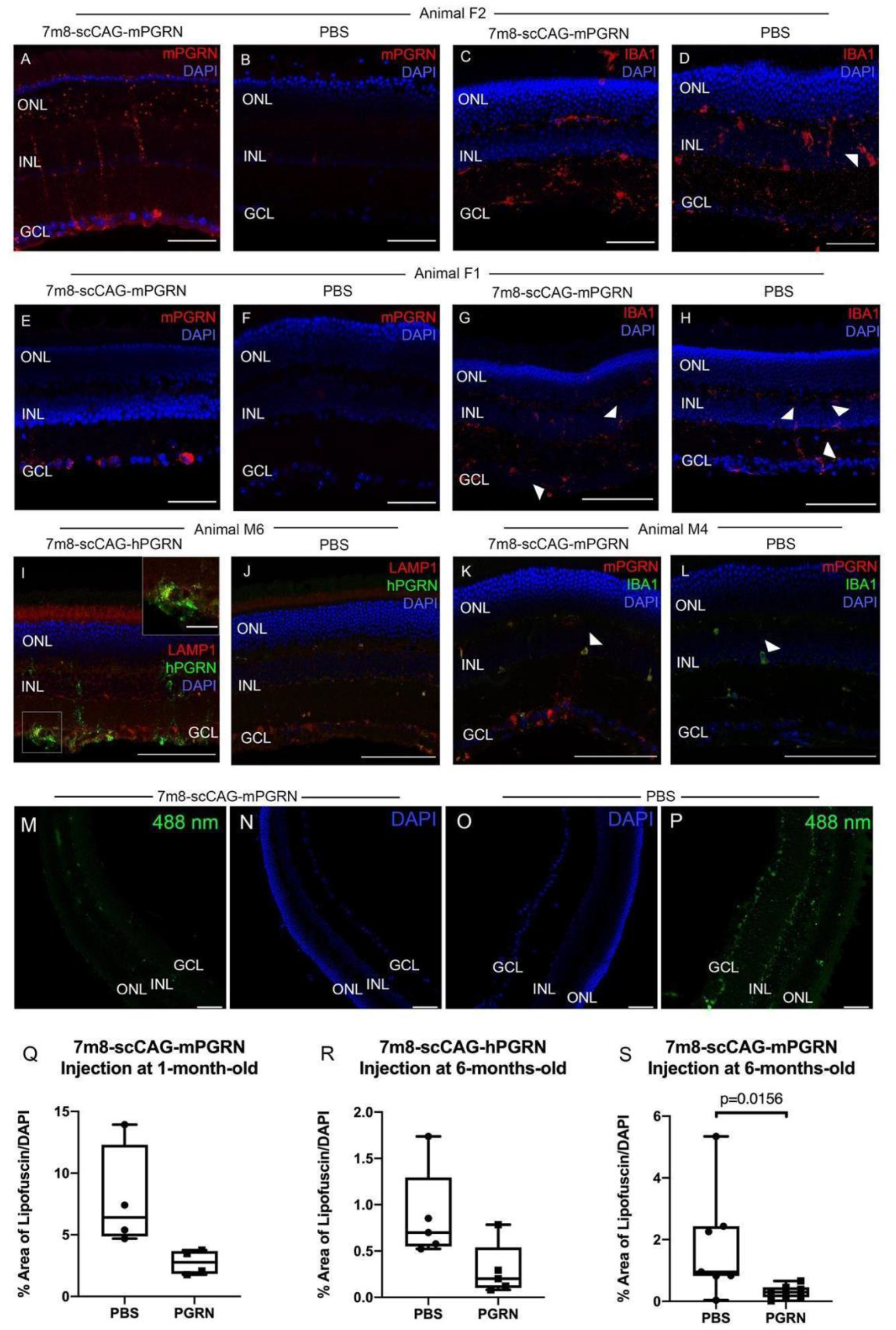
Immunohistochemistry on retinas of PGRN^-/-^ mice treated at 6 months of age, and injected with either 7m8-scCAG-mPGRN (A, C, E, G, K), 7m8-scCAG-hPGRN (I) or PBS (B, D, F, H, J, L). Mice show PGRN expression mainly in RGC, but also in bipolar cells, Müller cells and photoreceptors (A, E, I). IBA1 shows microglia migrating to INL and GCL (arrowheads) in all sections (D, G, H, K, L) except C. LAMP1 indicates little overlap between PGRN and lysosomes, with the exception of some puncta (inset – I). Similarly, there is little overlap between PGRN and microglia (K). DAPI (blue), mPGRN (red), hPGRN (green), IBA1 (red or green, as indicated). Lipofuscin can be seen autofluorescing in PGRN^-/-^ retina cross-sections under 488 nm-laser (M, N, O, P). Retinas expressing mouse PGRN visibly present a reduced amount and intensity of lipofuscin deposits (M). Quantification of lipofuscin deposits as the percentage of overall lipofuscin over total DAPI shows statistically significant reduction of lipofuscin in retinas transducing mouse PGRN at 6 months (S) but not at 1 month of age (Q) or expressing human PGRN at 6 months of age (R). Box and whiskers plot, boxes show interquartile range (25% to 75%), whiskers represent minimum and maximum values, horizontal lines represent median values of 4 animals (Q), 5 animals (R) and 7 animals (S). Wilcoxon signed rank test between eyes transducing progranulin and contralateral control eyes, p=0.0156. Lipofuscin (green) and DAPI (blue). Scale bars: 100 μm.

PGRN^-/-^ retinas expressing mPGRN showed a remarkable reduction in lipofuscin deposits (Figure 2S). Lipofuscin is visible in retina cross-sections, and the reduction in retinal lipofuscin in eyes transducing progranulin is easily observable when compared to levels in the contralateral control eyes (Figure 2M, P). Animals receiving intravitreal injections of mPGRN at 6 months of age showed a significant reduction in lipofuscin deposits when compared to contralateral eyes that received PBS injections (Figure 2S), but not at 1 month of age (Figure 2Q). On the other hand, animals transducing human PGRN showed a similar, but non-significant, reduction in retinal lipofuscin. (Figure 2R).

Despite improvements measured in terms of reduced retinal lipofuscin deposits there were no significant differences in retina thickness between treated and control eyes as quantified by retinal layer measurements through OCT imaging (Figure 3A, B, C). Retinal thickness was measured at 12 months of age from OCT images collected from animals injected at 1 and 6 months of age, and there were no differences between treated and control eye in total retina thickness or photoreceptor layer thickness, with the exception of one point in the inferior retina of animals injected with 7m8-scCAG-mPGRN at 6 months of age. There is a trend, albeit non-significant, towards reduction in total retina and photoreceptor layer thickness in treated eyes versus control eyes for animals injected at 1 month old (Figure 3A), as well as in total retina of animals injected at 6 months of age (Figure 3B, C). Analysis of ONL nuclei numbers corroborated these findings. Results for animals injected with 7m8-scCAG-mPGRN or 7m8-scCAG-hPGRN at 1 or 6 months of age and euthanized at 12 months of age are shown in Figure 3D-F. There was a significant reduction in photoreceptor nuclei in the retinas of animals injected with 7m8-scCAG-mPGRN at 6 months (Figure 3F), but not in the retinas of animals injected at 1 month of age (Figure 3D), or in retinas of animals injected with hPGRN at 6 months (Figure 3E).

**Figure 3.**
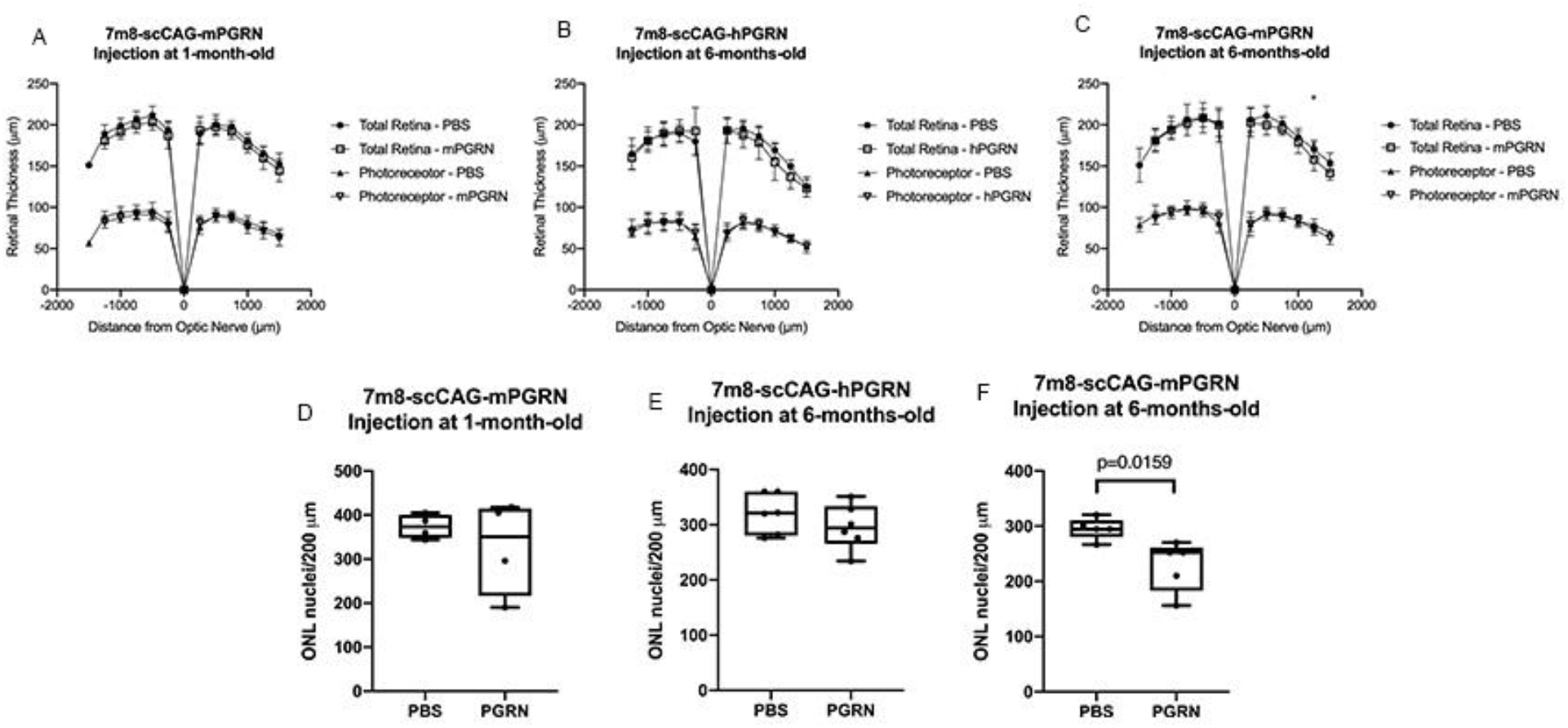
OCT measurements and ONL quantification in animals injected at 1 and 6 months of age. OCT quantification in superior (−250 to −1500 μm from the optic nerve) and inferior (250 to 1500 μm from optic nerve) PGRN^-/-^ retinas. Total retina thickness was measured from the ILM to RPE and photoreceptor layer thickness measured from ONL to RPE in PGRN^-/-^ mice intravitreally injected with 7m8-scCAG-mPGRN at 1 month of age (A), 7m8-scCAG-hPGRN at 6 months of age (B), or 7m8-scCAG-mPGRN at 6 months of age (C). Curves compare total thickness of retinas expressing PGRN (open square) and control retinas that received PBS (closed circle), or photoreceptor layer thickness of the same retinas expressing PGRN^-/-^ (open triangle) and control retinas that received PBS (closed triangle). Error bars represent ± standard deviation, and circles or squares represent mean values for retinal thickness for 10 animals per group in A and C, and 8 animals per group in B. Statistical tests were performed at each distance from optical nerve between mouse strains (t-tests), asterisk representing p=0.035. ONL nuclei in retina cross-sections were quantified over 200 μm sections in mice injected at 1 month of age (D), 6 months of age and expressing hPGRN (E), or 6 months of age and expressing mPGRN (F). Box and whiskers plot, boxes show interquartile range (25% to 75%), whiskers represent minimum and maximum values, horizontal lines represent median values of 5 animals. Mann-Whitney statistical test between control and treated retinas, p=0.0159, N=4 (D), N=6 (E) and N=5 (F).

Mice intravitreally injected at 12-months-old with 7m8-scCAG-mPGRN were euthanized at 18-months-old, enucleated and retinal tissue collected for processing. Immunohistochemistry showed the presence of transduced mPGRN in eyes that received 7m8-scCAG-mPGRN, but not PBS controls, as expected (Figure 4A, B, C, E, F). Once again, PGRN expression was highest in the RGCs, but was also expressed in bipolar cells, Müller glia and photoreceptors (Figure 4A, C, E). Once again there was large variability in expression patterns, despite being injected with the same viral stock (2 μl at 3.97E+13 vg/ml). Similar to Figure 2, IBA1 labelling showed microglia infiltrating mainly the INL, but also the GCL and ONL, in both treated and control retinas. However, treated retinas showed fewer microglia infiltration events than control ones (arrows, Figures 4G and H). Interestingly, the retinas of mice expressing mPGRN were thinner than those injected with PBS (Figure 4I). This statistically significant difference was observed in total retinal thickness, and the thickness of the photoreceptor layer at the inferior retina. Similar to the previous cohort of animals injected with 7m8-scCAG-mPGRN at 6 months of age (Figure 2), PGRN^-/-^ retinas expressing virally delivered PGRN showed a dramatic and statistically significant decrease in lipofuscin deposits in comparison to the controls injected with PBS (Figure 4J). The trend in ONL nuclei numbers was also comparable to previous results in animals injected at 1 and 6 months of age, whose retinas expressing PGRN showed a non-statistically significant trend of a reduction in photoreceptor nuclei in comparison to controls injected with PBS (Figure 4K).

**Figure 4.**
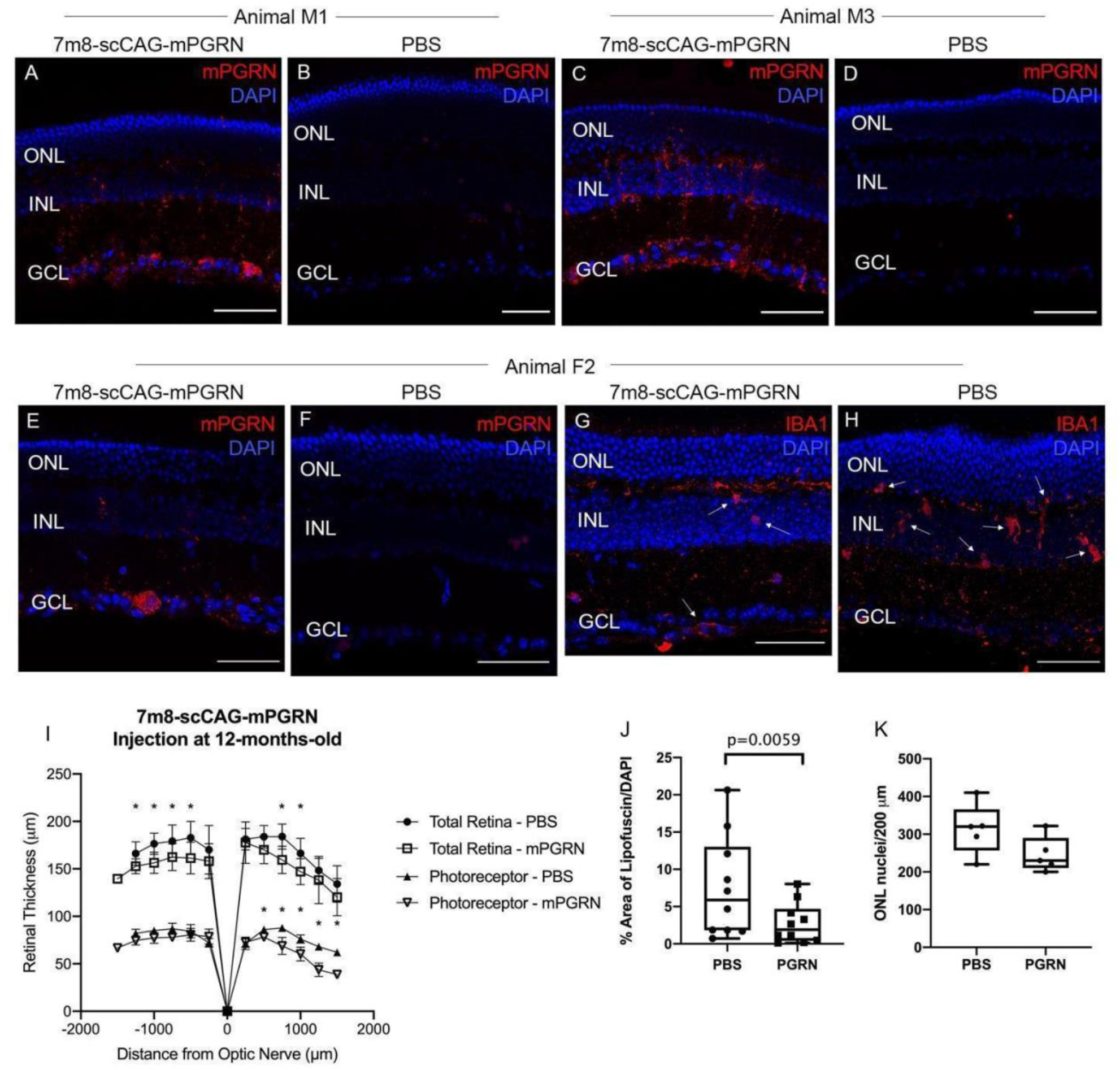
Immunohistochemistry on retinas of PGRN^-/-^ mice injected at 12 months of age with 7m8-scCAG-mPGRN (A, C, E, G) or PBS (B, D, F, H). Mice show mPGRN expression mainly in RGC, but also in bipolar cells, Müller cells and photoreceptors (A, C, E). IBA1, a microglia marker, shows microglia migrating to INL and GCL (arrows) in both treated and control retinas (G, H). However, microglia seem to be infiltrating nuclear layers more frequently in control mouse (H). DAPI (blue), mPGRN (red), IBA1 (red). Scale bars: 50 μm. OCT measurements for superior (−250 to −1500 μm from the optic nerve) and inferior (250 to 1500 μm from optic nerve) PGRN^-/-^ retinas (I). Total retina thickness was measured from the ILM to RPE and photoreceptor layer thickness measured from ONL to RPE in PGRN^-/-^ mice, showing retinal thinning in mice transducing PGRN. Curves compare total thickness of retinas expressing PGRN (open squares) and control retinas that received PBS (closed circles), or photoreceptor layer thickness of the same retinas expressing PGRN (open triangles) and control retinas that received PBS (closed triangles). Error bars represent ± standard deviation, and circles, squares or triangles represent mean values for retinal thickness of 8 animals per group. Statistical tests were performed at each distance from optical nerve between mouse strains (t-tests), asterisks representing p < 0.05. Quantification of lipofuscin deposits as the percentage of overall lipofuscin over total DAPI shows statistically significant reduction in lipofuscin in retinas transducing mouse PGRN (J). ONL nuclei in retina cross-sections were quantified along 200 μm sections for treated and control retinas (K). Box and whiskers plot, boxes show interquartile range (25% to 75%), whiskers represent minimum and maximum values, horizontal lines represent median values. Wilcoxon signed rank test between control and treated retinas, p=0.0059 and p=0.125 (ns) for N=10 (J) and N=5 (K), respectively.

Mouse pups injected intravenously at post-natal day 3 or 4 (P3-4) were euthanized at 12-months-old, perfused and enucleated. Immunohistochemistry for mPGRN and hPGRN showed an increase in PGRN expression in PGRN^-/-^ mice injected with AAV9.2YF-scCAG-mPGRN and AAV9.2YF-scCAG-hPGRN, respectively (Figure 5A, C). Once again, progranulin was mainly observed in RGCs (Figure 5A), although also found in bipolar cells and Müller glia (Figure 5C). Retinas from naïve control animals showed no PGRN expression (Figure 5B).

**Figure 5.**
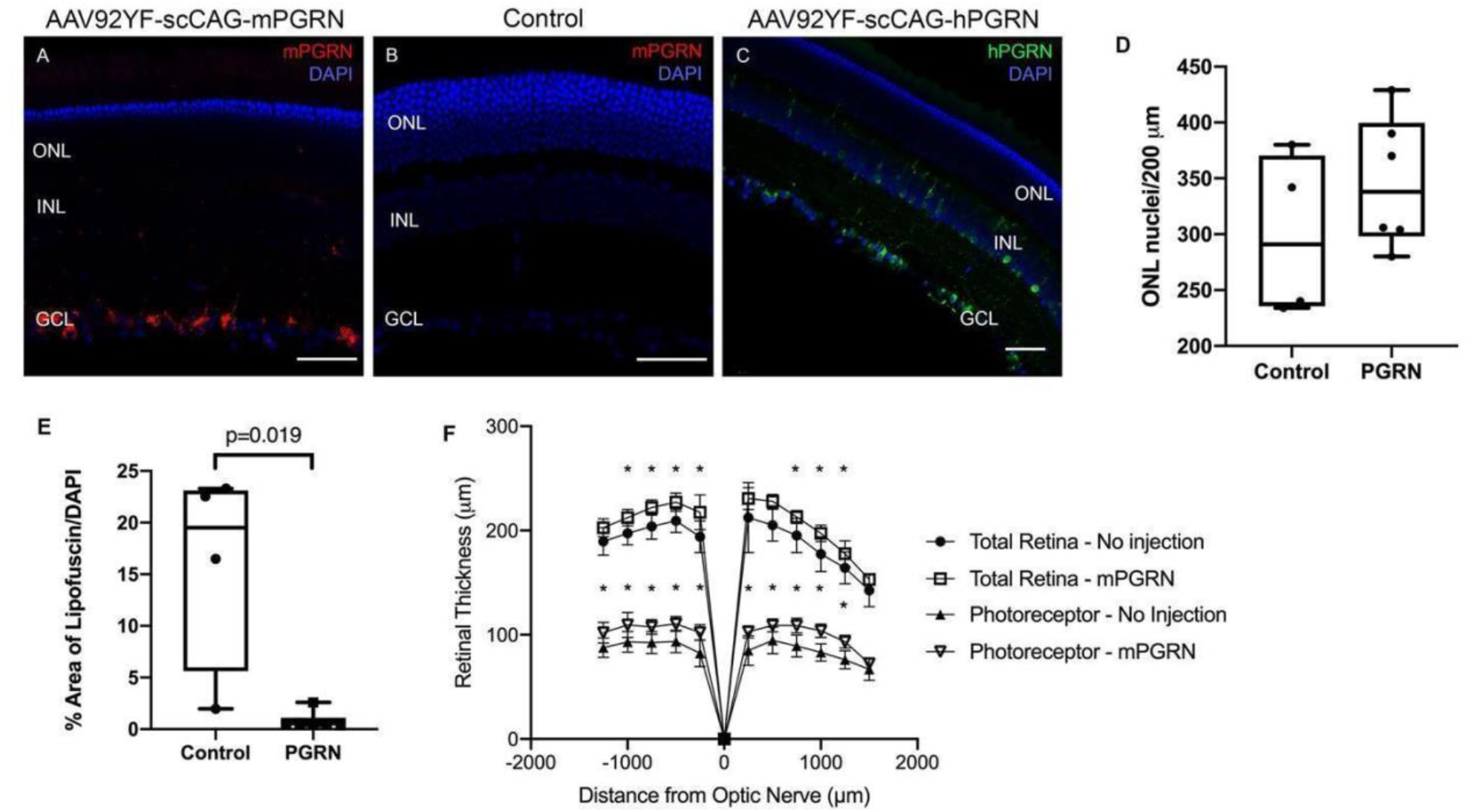
PGRN expression, ONL nuclei and lipofuscin area quantification. Immunohistochemistry for PGRN in retinas of mice intravenously injected with AAV92YF-scCAG-mPGRN (A), AAV92YF-scCAG-hPGRN (C) or not injected (B). DAPI (blue), mPGRN (red), hPGRN (green). Scale bars represent 50 μm (A-B) and 100 μm (C). ONL nuclei in retina cross-sections were quantified over 200 μm sections for treated and control retinas (D). Quantification of lipofuscin deposits as the percentage of overall lipofuscin over total DAPI shows statistically significant reduction in lipofuscin in retinas transducing mouse PGRN (E). Box and whiskers plot, boxes show interquartile range (25% to 75%), whiskers represent minimum and maximum values, horizontal lines represent median values. Mann-Whitney statistical test between control and treated retinas, p=0.352 (ns) (D) and p=0.019 (E) (N=4 (PBS) and N=6 (PGRN)). OCT measurements for superior (−250 to −1500 μm from the optic nerve) and inferior (250 to 1500 μm from optic nerve) PGRN^-/-^ retinas (E). Total retina thickness was measured at 12 months of age, from the ILM to RPE, and photoreceptor layer thickness measured from ONL to RPE in PGRN^-/-^ mice intravenously injected with AAV92YF-scCAG-mPGRN at post-natal day 3 or 4 (P3-4). Curves compare total thickness of retinas expressing PGRN (open squares) and naïve control retinas (closed circles), or photoreceptor layer thickness of the same retinas expressing PGRN (open triangles) and control retinas (closed triangles). Error bars represent ± standard deviation, and circles or squares represent mean values for retinal thickness of 6 animals per group. Statistical tests were performed at each distance from optical nerve between mouse strains (t-tests); asterisks represent p < 0.05.

Curiously, there was a trend in detecting more photoreceptor nuclei in mice injected with AAV9.2YF-scCAG-mPGRN than in control mice (Figure 5D). Although not statistically significant, this trend goes against what has been reported in Figures 2 and 4, where mice intravitreously injected with 7m8-scCAG-PGRN showed a trend of photoreceptor loss and retina thinning. However, the significant reduction in lipofuscin deposits was seen in these mice as well, indicating that despite the difference in photoreceptor numbers, PGRN still prevented lipofuscin accumulation (Figure 5E).

OCT imaging revealed a significant increase in total retina thickness of mice transducing mPGRN, and a significant overall increase in the photoreceptor layer as well (Figure 5F). In this case too, these trends are opposite to the trends reported for animals injected at 1, 6 and 12 months of age, which showed reductions in total retinal thickness when PGRN was expressed (Figure 3 and 4). In the mice injected at P3-4, the opposite was observed, with increased retinal thickness when compared to naïve PGRN^-/-^ littermates.

Moreover, when looking at the hippocampus of these mice there was also a striking difference in lipofuscin deposits (Supplemental Figure 2). Brain sections of both injected and naïve control mice were compared by imaging the dentate gyrus (DG), field CA1 and third ventricle (V3). Lipofuscin reduction in four mice expressing PGRN corroborated progranulin’s ability to reduce lipofuscin accumulation, when compared to four naïve controls (Supplemental Figure 2E).

## Discussion

The hypothesis driving the current study was that by delivering progranulin to PGRN^-/-^ mice it would be possible to prevent or slow down retinal degeneration, with earlier intravitreal injections having greater therapeutic effect. However, we observed something altogether more complex: earlier gene delivery in adult mice did not halt or prevent retinal degeneration, with delivery in elderly animals leading to further tissue loss. On the other hand, consistent reduction in lipofuscinosis and microglia infiltration indicated some level of improvement. Nevertheless, delivery to mouse pups elicited an improvement in both retinal thickness and lipofuscinosis.

Accumulation of autofluorescent lipofuscin represents a reported indicator of NCL disease, both in the brain and retina, and can be detected by fluorescence microscopy(14). It was a quantifiable biomarker for CLN11 and other NCL models in previous studies, and as such, establishing the levels of lipofuscin deposits in wild-type and knockout retinas over time was important. Initial experiments characterizing the model showed that lipofuscin could be used as a reliable biomarker of the disease process, and Ward and colleagues report that lipofuscinosis increases in PGRN^-/-^ mice are statistically significant as early as 1.5 months old in comparison to C57BL/6J controls(34). However, in this study, deposits are significantly increased in PGRN^-/-^ mice in comparison to C57BL/6J only at 12 and 18 months of age (Figure 1E). The difference can potentially be explained due to quantification methods: Ward and colleagues quantified lipofuscin puncta per mm^2^, while in this study the area of lipofuscin was quantified and normalized to DAPI. Similarly, OCT results indicated months 12 and 18 as reliable endpoints for retinal thickness assessment due to significant differences between C57BL/6J and PGRN^-/-^ mice (Figure 1A-D). Similarly, by quantifying nuclei in the outer nuclear layer (ONL) it was possible to observe a statistically significant reduction in photoreceptors in the ONL of PGRN^-/-^ mice when compared to C57BL/6J retinas at 12 months of age (Figure 1I). Retinal degeneration is slow in this model, and there is no complete loss of photoreceptors; cones and rods obscured B wave results in ERGs, where only scotopic A waves showed differences between mouse strains at 7, 9 and 12 months of age (Supplemental Figure 1). These response patterns are consistent with histological data showing relatively little photoreceptor cell death in PGRN^-/-^ mice over the first 6 months but a significant reduction in the overall number of photoreceptors by 12 months of age. Similarly, as rods are the most abundant photoreceptor in the mouse retina, it is expected that scotopic ERG recordings would be more sensitive to the loss of photoreceptors. Therefore, ERG comparisons were not considered a reliable indicator of disease progression for use in evaluating therapeutic efficacy, but rather served to confirm the results seen in OCT imaging on an electrophysiological level.

The AAV2.7m8 capsid was chosen for intravitreal injections in adult mice due to its ability to infect cells in all layers of the retina from the vitreous. Since loss of progranulin leads to loss of photoreceptors and thinning of the ILM and GCL in human patients and knockout mice, it was preferable to target all retinal layers(21). Furthermore, similar to the AAV2 and AAV5 vectors, intravitreal delivery of 7m8 causes a humoral immune response that prevents further transduction after readministration but does not cause tissue damage due to immune response in mice(30). The AAV9.2YF capsid was chosen for intravenous delivery in mouse pups due to its ability to cross the blood-retina and blood-brain barriers in early postnatal days, therefore delivering the viral load to both tissues, and before an intravitreal delivery would be possible.

Reduction in lipofuscin deposits was a consistent trend in PGRN^-/-^ mice that received either 7m8-scCAG-PGRN or AAV9.2YF-scCAG-PGRN (Figure Q-S, Figure 4J, Figure 5E). While it was expected that mice expressing PGRN from young adulthood might show robust reduction in overall lipofuscin accumulation, the opposite was observed. Mice injected at 6 or 12 months of age showed the most dramatic reductions in lipofuscin accumulation. Interestingly, mice intravenously injected at P3-4 had similar retinal reductions in lipofuscin as mice intravitreously injected at 6-months-old (p=0.0190 and p=0.0156, respectively), while mice treated at 12 months of age showed the most dramatic reduction among all 4 groups (p=0.0059). However, it must be noted that mice treated at 12 months of age were euthanized at 18 months of age, 6 months later than animals in the other groups. This extended time frame for accumulation of lipofuscin deposits might have exaggerated the differences between treated and control eyes. Finally, the treatment also apparently reduced lipofuscin accumulation in the brains of mice intravenously injected with AAV9.2YF-scCAG-mPGRN (Supplemental Figure 2).

The retinal thickness data obtained by OCT imaging tell a different story. Here we can see the critical role that OCT imaging can play in allowing quantification of retinal changes in live animals at different timepoints; it also circumvents the problem of histological artifacts in quantification. While cohorts treated at 1 and 6 months of age showed a slight trend in retinal thinning when expressing progranulin in comparison to the control eye (Figure 3A-C), only animals injected at 12 months of age exhibited statistically significant reductions in retinal thickness (Figure 4I). This suggests that progranulin, while potentially improving lysosomal function and reducing lipofuscin deposits, may exacerbate the loss of photoreceptors. However, once again it must be pointed out that the first two groups treated at 1 and 6 months of age were euthanized at 12 months of age, while the third cohort was euthanized at 18 months of age. Thus, it is possible that the more exaggerated changes in the latter group are a product of further aging rather than a reflection of the age of treatment. Additional experiments are needed to distinguish between these possibilities.

The only group to bear out the proposed hypothesis that transducing PGRN would slow retinal degeneration was the group injected at P3-4, showing significant improvement in retinal thickness when transducing progranulin (Figure 5F). The result suggests that earlier treatment not only leads to improvement in lysosomal function, but also reduces photoreceptor cell death. If this is due to early or systemic delivery, or a combination thereof, is unknown. However, the observation that this improvement was only present when viral delivery was made at a very early, neonatal timepoint raises the question of the translatability of the results to the clinical scenario. Nevertheless, the results do represent a proof-of-concept that progranulin delivery and expression can rescue retinal cells.

Interestingly, immunohistochemistry showed the variability of PGRN expression in retinas of different littermates injected by the same surgeon and with the same viral aliquot. It is also important to note that immunolabeling reported in this study showed intracellular progranulin and does not represent any progranulin that was potentially secreted by infected cells. Furthermore, immunohistochemistry results showed fewer microglial infiltration events in retinas transducing PGRN after injections at 6 months of age (Figure 2C, D, G, H, K, L). These results corroborated previous reports of increases in activated microglia in the hippocampus of PGRN^-/-^ mice (35) and indicated that delivery of PGRN potentially minimizes microglial activation in the retina. Further exploration of microglial activation in retinas expressing PGRN is warranted.

Moreover, human PGRN does not seem to colocalize with LAMP1, a lysosomal transmembrane protein and marker, with the exception of limited puncta in retinal ganglion cells (Figure 2I). The question of correct lysosomal localization of transduced PGRN in neurons remains unanswered. Additionally, it is important to note that hPGRN did not prove toxic when delivered to PGRN^-/-^ mice either intravitreally or intravenously.

In the 2018 study by Arrant and colleagues, PGRN^-/-^ mice were stereotaxically injected rAAV2-CBA-mPGRN in the mPFC and showed reduction in microgliosis and lipofuscinosis in different brain regions(27). They demonstrated an improvement in lysosomal function by the analysis of cathepsin D expression and LAMP1. Most curiously, they also indicated that PGRN^-/-^ mice mount a strong local immune response to transduced PGRN, associated with a higher antibody production than compared to mice injected with rAAV2-CBA-GFP.

In a later study, Amado and colleagues (2019) examined histologically the brains of PGRN^-/-^ mice transducing progranulin(28). Stereotaxic injections of rAAV4-CAG-hPGRN or rAAV9-CMV-hPGRN in the posterior right lateral ventricle led to atrophy of the hippocampus in PGRN^-/-^ mice, linked to toxic CD4 T-cell infiltration. Interestingly, animals injected with AAV9-CMV-GFP or AAV4-CAG-GFP did not show the same level of hippocampal damage or cellular infiltrate as AAV9-CMV-hPGRN or AAV4-CAG-hPGRN, indicating that degeneration occurred due to transduction of progranulin and not as a response to the viral capsid. They also reported an increase in microgliosis and astrogliosis in the hemisphere transducing PGRN, and even an inflammatory response in wild-type mice injected with AAV and over-expressing PGRN.

The retinal changes in response to progranulin expression observed in our study seem to fall in line with observations in the Arrant and Amado studies, as we also observed a strong reduction in lipofuscinosis, and an apparent reduction in microgliosis. However, despite these positive findings, there was a noticeable reduction in retina thickness and photoreceptor nuclei in injected adult animals. That animals injected at P3-4 did not exhibit retinal thinning, suggesting that this adverse treatment effect is time-sensitive. It is also important to note the differences in cell targeting by AAV2.7m8 and AAV9.2YF, wherein the former only offers PGRN transduction in the retina and the latter results in systemic expression of PGRN. An immune response to PGRN is offered as a possible explanation. PGRN^-/-^ mice are known to present an exacerbated immune response to injury or stressors, which can be a contributing factor to the results observed(35–37). PGRN is normally secreted by microglia and while it has not been confirmed that neurons and Müller glia transducing PGRN release it to the extracellular space, it is possible that PGRN^-/-^ mice exposed to this extracellular protein for the first time later in life mount an immune response to it. The delivery of PGRN to P3-4 mice is consistent with this explanation, as mice exposed to the protein from an early age might not mount an immune response to the protein. More comprehensive studies of systemically treated PGRN^-/-^ pups are warranted, as well as studies to specifically investigate the possibility of an immune response against PGRN in animals treated at an older age. It is also still unknown if the transduced PGRN, delivered via AAV, leads to correct protein shuttling and signalling within the cell. While immunohistochemistry for PGRN in our study points towards vesicular localization, this finding requires confirmation. Nevertheless, the positive outcomes for mice injected at an early age, such as reduction in lipofuscinosis, microgliosis and increase in retinal thickness, are promising. The effect of PGRN gene therapy in the retina serves as a proof of point for further investigating PGRN gene therapy in PGRN-related NCL disease.

## Acknowledgments

The research was supported by the Foundation Fighting Blindness grant TA-GT-0818-0745-UCB. The authors would like to thank Michael Ward and Ari Green at UCSF for supplying the original mouse breeding pairs, Mei Li for helping with viral preparations, Cameron Baker with manuscript editing, and the CNR Biological Imaging Facility, UC Berkeley. Research reported in this publication was supported in part by the National Institutes of Health S10 program under award number 1S10RR026866-01. The content is solely the responsibility of the authors and does not necessarily represent the official views of the National Institutes of Health.

## Author Contributions

Conceived and designed the experiments: EAZ, MK, JGF. Performed the experiments: EAZ, MV, MK, DH, JT, NMW. Analyzed the data: EAZ, DH, JT, NMW, JGF. Wrote the manuscript: EAZ, JGF.

## Supplemental Figures

**Supplemental Figure 1.**
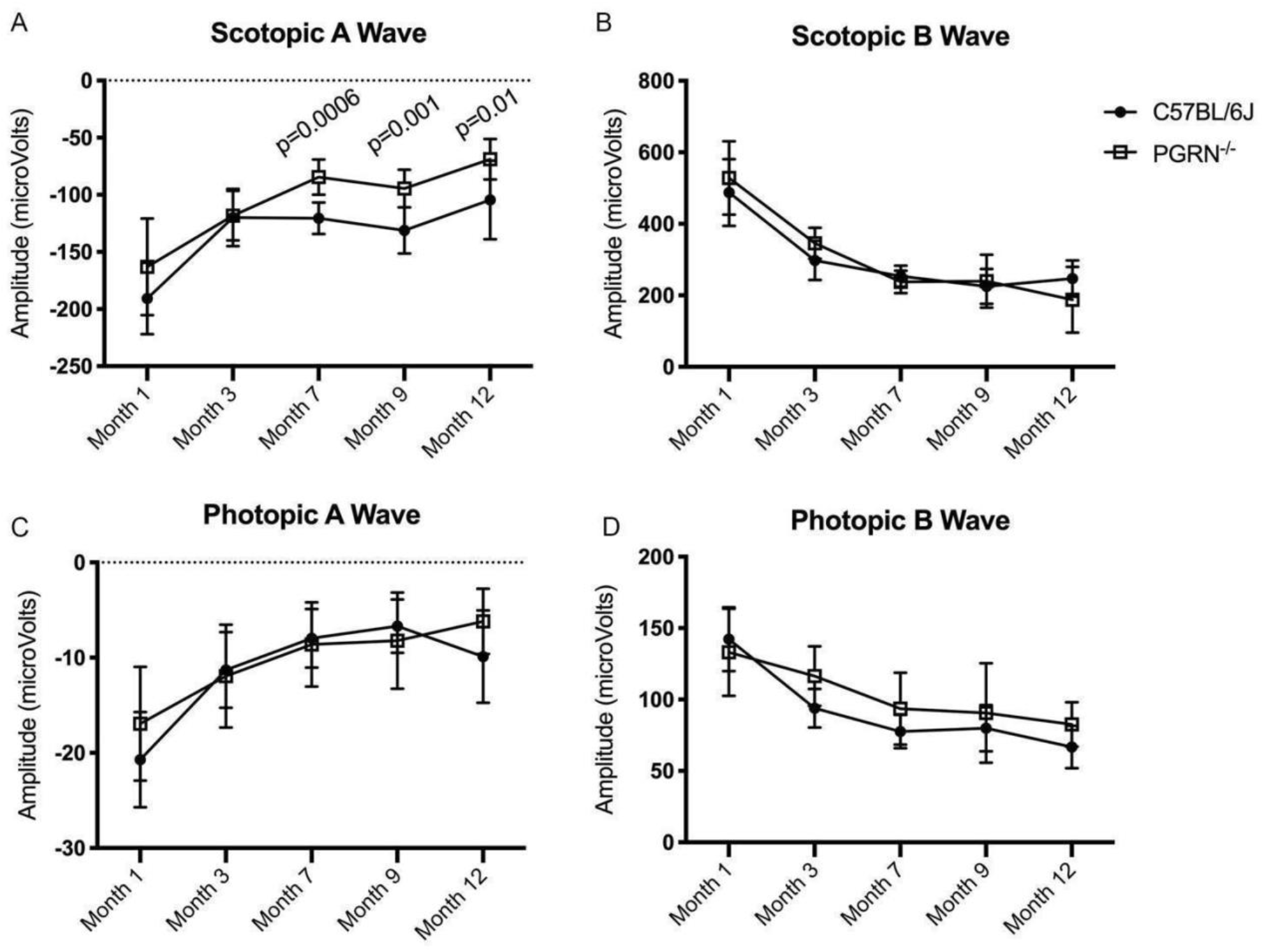
Electroretinogram (ERG) recordings from age-matched C57BL/6J and PGRN^-/-^ mice. Mean A wave and B wave amplitudes derived from scotopic (A, B) and photopic (C, D) recordings in C57BL/6J (closed circles) and PGRN^-/-^ (open squares) mice, plotted against animal age. Differences between strains were significant only at months 7, 9 and 12 for scotopic A-wave amplitude (Mann-Whitney statistical test; p values as indicated). Error bars represent ± standard deviation, circles and squares represent the mean value, 8 animals per group.

**Supplemental Figure 2.**
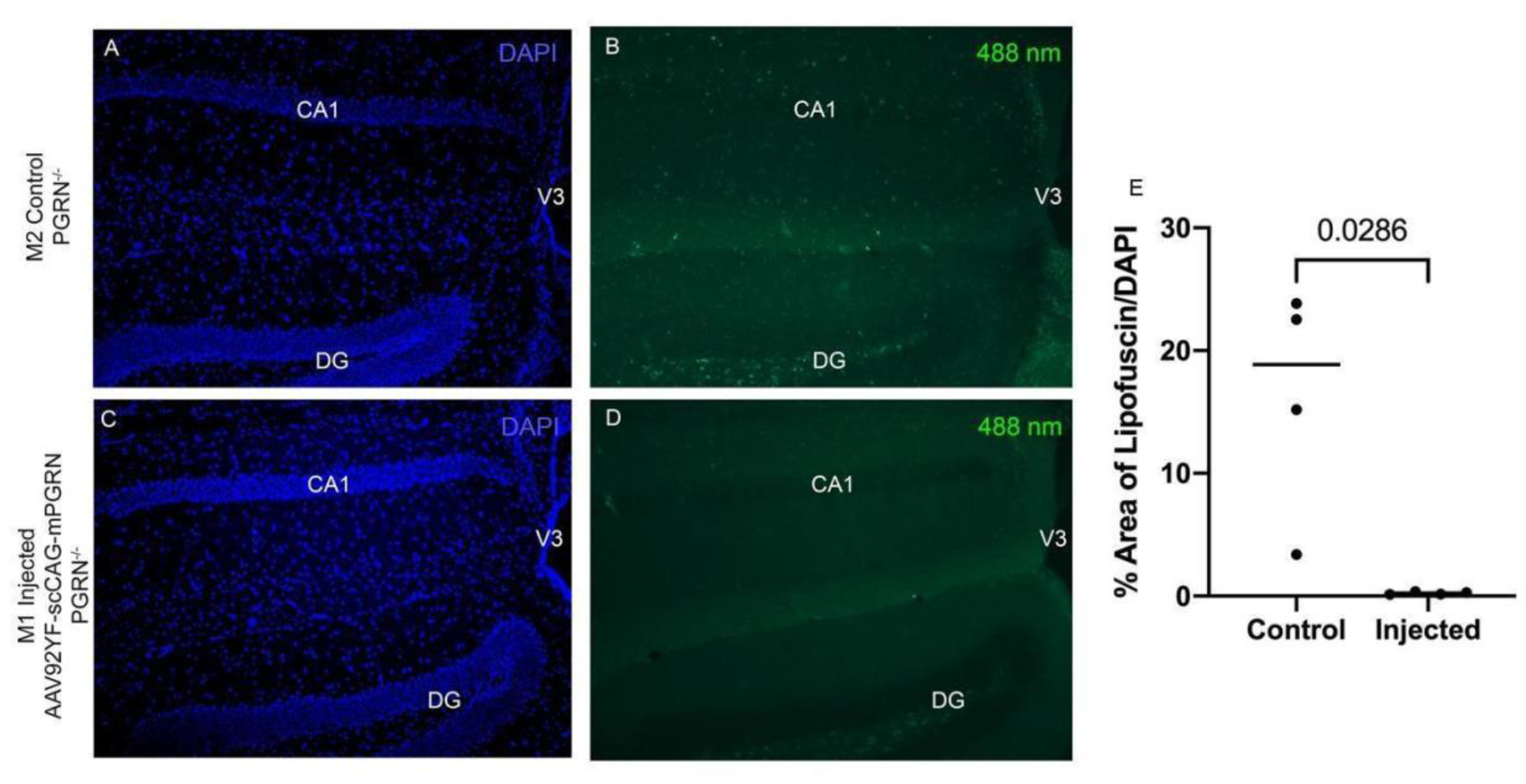
Lipofuscin deposits in the hippocampus. Coronal sections of mouse hippocampus focusing on the dentate gyrus (DG), field CA1 and third ventricle (V3) in control (A-B) and AAV92YF-scCAG-mPGRN injected mice (C-D) show accumulation of lipofuscin. Lipofuscin deposits, expressed as the percentage of overall lipofuscin over total DAPI, show reduction of lipofuscin in hippocampi transducing mouse PGRN; lipofuscin is reduced in hippocampus of mouse transducing PGRN (E). Plot shows individual values, with horizontal line representing median. Mann-Whitney statistical test between control and treated brain sections (p=0.0286) and N=4 for each experimental group. DAPI (blue) and autofluorescent lipofuscin (green).

